# Differences in selective profiles among single or double stranded DNA or RNA viruses. Statistical detection of its double or single stranded condition

**DOI:** 10.1101/2024.01.03.574038

**Authors:** Carlos Y Valenzuela

## Abstract

I developed a test to study the probability distribution of bases in dinucleotides separated by 0, 1, 2…K nucleotides sites. This analysis define selective profiles of dinucleotides by their sign and selection coefficients, their distance to neutrality measured by a chi-square value and the order of significance of their distance to randomness among the 16 possible values. In double stranded (ds) DNA or RNA an index (index = I, with 5’-3’ sense) dinucleotide participates in evolutionary processes together with three dinucleotides defined by base complementarity and 5’-3’ sense. The anti-sense dinucleotide in the same strand, the parallel (Par, with 3’-5’ sense) and anti-parallel (a-Par, with 5’-3’ sense) dinucleotides in the complementary strand. Single stranded (ss) RNA or DNA do not have these set of dinucleotides and would not present similarities or differences, among those four dinucleotides. I studied viromes taken from GenBank, with ds and ss RNA and DNA by this dinucleotide analysis. The core matrix has rows with Separations (Sep) of bases and columns with their selective profiles of the Index, Par and a-Par dinucleotides. Profiles distances I-Par and I-a-Par where compared. The ds RNA or DNA showed larger selective profiles differences between I-Par than I-a-Par. Ss RNA or DNA viruses showed non-significant differences between these comparisons. Those ss virus that exist as ds virus in their hosts, as Hindbis and HIV virus behaved similarly to ds viruses. Coli phage PhiX174 an ss DNA virus showed smaller I-Par than I-a-Par selective distances.

**IMPORTANCE:** I present a new method for the comparison of selective profiles that allows know whether a RNA or DNA behaves preferentially as ds or ss nucleic acid and gives information about their dynamic or evolutionary mechanisms of replication cycles. This changes our vision of viruses constituted in the virion as ds or ss viruses towards a functional evolutionary vision where viruses are ds and ss according to their behavior along with their life cycle. This evolutionary vision may be crucial in epidemiology.

## INTRODUCTION

Searching for a measure of the distance to neutrality of genomes, I developed a test, based on the distance to neutrality of dinucleotides whose bases are separated by 0, 1, 2,…K nucleotide sites (1–5). I used as the measure of the distance to neutrality a chi-square (χ^2^) test of the random distribution of the first base according to the second base of the dinucleotides (1–5). This study produces a matrix where rows are the number of nucleotide sites between both bases (Separations = Sep) and whose columns are the order of the distance to neutrality the 16 dinucleotides have. This order follows the value of the χ^2^ test assigned to a dinucleotide among the 16 values and it may or may not include the sign of the selection coefficient to make more or less discriminant the test. There is a total χ^2^ value given by the addition of the particular χ^2^ value for each dinucleotide. The particular χ^2^ value has one degree of freedom (χ^2^_1_); the total χ^2^ value has nine degrees of freedom (χ^2^_9_), because it correspond to the 4×4 matrix (4 rows x 4 columns, for the four bases) that loses one degree of freedom due to fixed marginal (rows or columns) totals. Distances to neutrality resulted enormous. The significance at the 0.05 level for χ^2^_1_ corresponds to 3.84; I found 377.3 (see Table 1) in *Sars-Cov-2* virus, for dinucleotide CG, with separation 0 (contiguous bases); the probability (P) is out of tables and programs; with Gaussian approximation (5), P < 10^-50^. The significance at the 0.05 level for χ^2^_9_ corresponds to 16.9; I found in human chromosome 21 χ^2^_9_ = 1,885,266.8 (5) for Sep 0 (P < 10^-222,180^).

**Table 1.** Significance vs Separations of dinucleotides from *SARS-CoV-2*. Four Separations.

| Sep | 1° Significance |  |  | 2° Significance |  |  | 3° Significance |  |  | 4° Significance |  |  |
| --- | --- | --- | --- | --- | --- | --- | --- | --- | --- | --- | --- | --- |
| | dinuc[s] | $\chi^2_1$ | co-sel | dinuc[s] | $\chi^2_1$ | co-sel | dinuc[s] | $\chi^2_1$ | co-sel | dinuc[s] | $\chi^2_1$ | co-sel |
| 0 | CG[-] | 377.3 | -0.5921 | AT[-] | 110.0 | -0.1959 | TA[-] | 85.7 | -0.1728 | TC[-] | 69.0 | -0.1980 |
| 1 | GT[+] | 28.6 | 0.1232 | TC[+] | 8.0 | 0.0673 | AA[+] | 5.6 | 0.0459 | CA[+] | 3.0 | 0.0424 |
| 2 | GG[+] | 55.5 | 0.2197 | TT[+] | 21.2 | 0.0830 | CC[+] | 8.7 | 0.0927 | AG[+] | 1.7 | 0.0315 |
| 3 | TG[+] | 15.5 | 0.0907 | CT[+] | 6.9 | 0.0627 | GT[+] | 4.7 | 0.0498 | TA[+] | 2.0 | 0.0264 |
| Sep | 5° Significance |  |  | 6° Significance |  |  | 7° Significance |  |  | 8° Significance |  |  |
| | dinuc[s] | $\chi^2_1$ | co-sel | dinuc[s] | $\chi^2_1$ | co-sel | dinuc[s] | $\chi^2_1$ | co-sel | dinuc[s] | $\chi^2_1$ | co-sel |
| 0 | CC[-] | 15.0 | -0.1218 | GA[-] | 11.4 | -0.0805 | GG[-] | 3.0 | -0.0508 | AG[-] | 0.098 | -0.0075 |
| 1 | AG[+] | 1.1 | -0.0168 | TG[+] | 0.47 | 0.0158 | CG[+] | 0.0 | 0.0053 | AC[-] | 0.079 | -0.0069 |
| 2 | AA[+] | 0.597 | 0.0150 | CA[+] | 0.189 | 0.0107 | TC[+] | 0.0 | 0.0044 | TA[-] | 0.039 | -0.0037 |
| 3 | GC[+] | 1.6 | 0.0389 | AG[+] | 1.5 | 0.0293 | AC[+] | 0.2 | 0.0099 | CC[+] | 0.011 | 0.0033 |
| Sep | 9° Significance |  |  | 10° Significance |  |  | 11° Significance |  |  | 12° Significance |  |  |
| | dinuc[s] | $\chi^2_1$ | co-sel | dinuc[s] | $\chi^2_1$ | co-sel | dinuc[s] | $\chi^2_1$ | co-sel | dinuc[s] | $\chi^2_1$ | co-sel |
| 0 | TT[+] | 5.96 | 0.0440 | GT[+] | 6.2 | 0.0573 | GC[+] | 7.8 | 0.0852 | AA[+] | 14.3 | 0.0730 |
| 1 | CT[-] | 0.372 | -0.0145 | TT[-] | 0.9 | -0.0168 | CC[-] | 2.5 | -0.0494 | GC[-] | 3.1 | -0.0532 |
| 2 | AC[-] | 0.32 | -0.0140 | GA[-] | 1.3 | 0.0150 | CT[-] | 1.8 | -0.0316 | AT[-] | 1.8 | -0.0252 |
| 3 | GA[-] | 0.014 | -0.0028 | AT[-] | 0.1 | -0.0066 | CA[-] | 0.3 | -0.0136 | AA[-] | 0.9 | -0.0136 |
| Sep | 13° Significance |  |  | 14° Significance |  |  | 15° Significance |  |  | 16° Significance |  |  |
| | dinuc[s] | $\chi^2_1$ | co-sel | dinuc[s] | $\chi^2_1$ | co-sel | dinuc[s] | $\chi^2_1$ | co-sel | dinuc[s] | $\chi^2_1$ | co-sel |
| 0 | CT[+] | 57.6 | 0.1809 | AC[+] | 87.7 | 0.2312 | CA[+] | 118.8 | 0.2690 | TG[+] | 267.19 | 0.3770 |
| 1 | TA[-] | 3.2 | -0.0336 | GA[-] | 5.3 | -0.0547 | GG[-] | 5.4 | -0.0683 | AT[-] | 8.3 | -0.0539 |
| 2 | CG[-] | 2.8 | -0.0514 | GC[-] | 5.7 | -0.0726 | GT[-] | 8.6 | -0.0677 | TG[-] | 33.9 | -0.1342 |
| 3 | TC[-] | 2.1 | -0.0349 | CG[-] | 7.8 | -0.0849 | TT[-] | 11.1 | -0.0601 | GG[-] | 14.8 | -0.1136 |

Sep = n° sites of separation; dinuc[s] = dinucleotide with its sign of selection; $\chi^2_1$ = chi-square value for the dinucleotide at the left hand; co-sel = selection coefficient.

Unexpectedly I found periodicity with period 3 in the χ^2^ value of distance to neutrality in virus and prokaryotes and period 2 and 6 in human chromosomes. This periodicity of the distance to neutrality is incompatible with neutral and nearly neutral evolution (2–5).

Selective values allow the characterization of selective profiles of dinucleotides (hereafter, pair and dinucleotide will be synonymous unless I state other precision). The selective profile of a dinucleotide includes: 1) its selective value, 2) the sign of this selection value, 3) its χ^2^ value and 4) the order of significance of this value (1° to 16°). The χ^2^ value is calculated by (Obs_i_ – Exp_i_)^2^/Exp_i_, where Obs_i_ and Exp_i_ are the observed and expected numbers of the i_th_ dinucleotide, respectively. The total χ^2^_9_ value is the sum of the 16 values of the 16 nucleotides. The selection coefficient and its sign is the quotient (Obs_i_ – Exp_i_)/Exp_i_, where positive coefficients (selectively advantageous) apply to dinucleotides with more observed than expected pairs and negative coefficients (selectively disadvantageous) apply to dinucleotides with less observed than expected pairs. Any dinucleotide has its selective profile and this profile (a profile of an index dinucleotide or I profile) compared with the other 15 profiles produces 15 distances (to the I profile).

In double stranded (ds) RNA or DNA mutation or selection (and eventually genetic drift) of one or the two bases of one dinucleotide occur synchronously among the four dinucleotides that a pair implies due to base complementariness and 5’-3’ sense involved in duplication or transcription processes. This does not happen in a single stranded (ss) DNA or RNA. Thus, the test of comparison among selective profiles can discriminate whether the nucleic acid of the organism in study is ds or ss DNA or RNA. In ds DNA as in the human genome, the Index (I) 5’G…A3’ dinucleotide has a contra-sense (C-S) pair 3’A…G5’ (in the same DNA strand), a parallel (Par) pair 3’C…T5’, and an anti-Parallel (a-Par) pair 5’T…C3’ pair in the complementary strand. These four dinucleotides evolve together and synchronously. This coordinated evolution is not possible (physically but not evolutionarily during the viral life cycle) in ss viruses. Since there is only one published strand (the index strand from GenBank) of the nucleic acid, I study the selective profile of these four dinucleotides in the index strand (all pairs in the sense 5’…3’) and determine the similitude or difference among their selective profiles. Regardless (a priori) the reproductive processes of the organism, if there are evolutionary differences in replication or transcription between the senses of the copy, or differences due to the specific pair, the test will detect these differences. In ss DNA or RNA, these differences should be impossible because there are not a complementary strand to generate those differences.

This study is on RNA or DNA viruses, however, the differences I study are not on the ss or ds physical constitution of viruses well stablished from the infective viral particle (virion). This study deals with on the evolutionary constitution and dynamic behavior of viruses. Either ds or ss viruses have different ds or ss processes along their life cycles.

Mutation, selection, genetic drift, evolutionary contingencies occur at any stage of the reproductive or vital viral cycle with different intensities and in mono or bi-stranded constitutions. As for example, ss retroviruses having a retro-duplication of its nucleic acid and existence as a temperate or inserted virus in the DNA of the host may have rather ds (evolutionary) nucleic acid selective profile. A ds virus whose stage of mutation is mainly at the ss replication process may have ss selective profile (this is very improbable).

## GENOMES AND METHODS

Complete genomes taken from GenBank (search in PubMed, Nucleotide, FASTA). **Double stranded DNA**: Human adenovirus C strain KF268127.1 Human USA CL 42 1988 (35,931 bp); Human cytomegalovirus strain AD169 X17403 (229,354 bp); Escherichia phage 2 vB EcoM PhAPEC2 KF562341 (167,318 bp); His2 virus NC_007918.1 (16,067 bp). **Single stranded DNA**: Penaeus monodon hepandensovirus 4 NC_011545.2 (6,310 b); Escherichia phage phi-X174 NC_001422 (5,386 b); Phage M13 Enterobacteria NC_003287.2 (6,407 b). **Double stranded RNA**: Marmot picobirnavirus strain HT1 KY855428.1 (4,579 bp); Gentian Kobu virus NC_020252.1 (22,711 bp). **Single stranded RNA**: *SARS-Cov-2* LR757998.1 Wuhan (29,866 b); Sindbis virus NC_001547.1 (11.703; HIV MN691959.1 isolate ACH2-NFLMDA13 B1 (USA) (9,943 b). *E coli* genome strain AVS0967 NZ_CP124398 (5,097,505 bp). The number of nucleotides used in the analyses may be less than the number in GenBank because the base identification is not complete. I did not study positive and negative condition of the viral strand to restrict the analysis.

### Method

– The nucleotide sequence of a genome, chromosome or nucleic acid segment constitute a primary file obtained from GenBank.

– From this file the sets of dinucleotides whose bases are separated by 0, 1, 2,…K sites are constructed.

– Each of these sets yield a matrix where rows are the four bases A, T, G, and C for the first base (downstream) and columns are the four bases A, T, G, and C for the second base (upstream). This is a matrix of 4×4 for the 16 possible dinucleotides. For each of the 16 dinucleotides with bases separated by K sites a χ^2^ test describes the distance to neutrality.

The expected number of each dinucleotides (Exp) is calculated by (fB_u_)x(fB_d_)xN, where f is the observed frequency of B, a generic base (A, T, G or C), u and d denote upstream and downstream, respectively and N is the total number of observed (Obs) dinucleotides for this K. The χ^2^_1_ (one degree of freedom = df) values for each dinucleotide is [(Obs_i_-Exp_i_)^2^/ Exp_i_), i going from 1 to 16. The total χ^2^_9_, with 9 df is the sum of the 16 values plus a minimal value corresponding to the difference of the Obs number of each dinucleotide with the total. This specific difference is small for each dinucleotide and neglected in present calculations.

– The selection coefficient with its algebraic sign proceeds from ((Obs_i_-Exp_i_)/ Exp_i_)

– Data are presented in a matrix where the rows are K (Sep = separations) and columns are selective values ordered according to their distance to neutrality (χ^2^_1_) or order of significance (Sig), sign of the selection coefficient and the value of the selection coefficient (co-sel). I stopped K at 33 separations, including 0 Sep. These selective values constitutes the selective profile of this pair. I used only four Seps to short the analysis.

– The estimates of the values of the profile is directly the figures just described. From these estimates the difference between the order of significance, selection coefficient and the sign of selection between the parallel (Par) or the anti-Parallel (a-Par) and the Index dinucleotide follows by direct subtraction. I did not include the (χ^2^_1_) value in the estimation of selective differences because genomes differ in the number of dinucleotides, and this value depends greatly of this number, nor I included the profile of the contra-sense pair.

– In the present study separations go from 0 (contiguous bases) to 3 sites of separations.

– Study these traits in Table 1.

### Some ad hoc statistical considerations and tools (modified from (5))

I analyzed all the genome dinucleotides of an organism, so parameters are known, statistical tests are not necessary. I applied statistical tests to show the robustness of the analyses and sense of differences. The distance of a selective profile between two types of dinucleotides has mean (M), variance (V) and standard deviation (SD). For example, the variable distance of Par or a-Par dinucleotides to the Index (ID) dinucleotide. These parameters consider all the possible distances occurring with equal probability. In Significance (Sig) comparisons, only absolute (differences without + - signs) distances are used. An algebraic full demonstration of estimates of M, V and SD is out of the scope of this article (OSA). I offered an intuitive but complete calculation (C.Y. Valenzuela, submitted for publication). The expected mean difference in Sigs between the Index profile and Par or a-Par profiles is M = 5.667 and its SD is 3.6362. With these parameters, I tested the observed figures with one tailed t tests (with the used sample sizes it does not differ from a z test) and equal variances. The initial difference between selective coefficients (co-sel) is calculated with direct figures; this difference have a mean equal to 0 (the set of I dinucleotides is the same of the set of Par and a-Par dinucleotides). The SD must approximate that of a uniform distribution theoretically between −1.0 to +1.0 (limits of co-sels); but since selection is strongly dependent on evolutionary contingencies, I used the observed variance to construct the test because variances proceed from the same source. The definitive comparison test must be on absolute differences to avoid the difference between means that is 0, what implies distance equal to 0. The χ^2^_1_ value is always positive and hides the direction of selection. I ordered those values according to a cline from the most positively to the most negatively selected or vice versa. The previous analysis (C. Y. Valenzuela, submitted for publication) considered only the magnitude of the deviations to neutrality the present one includes the sense of selection; that is why the Table in reference (C. Y. Valenzuela, submitted for publication) is different with the present one. It is irrelevant for the order of dinucleotides whether they begin from the most negative or the most positive pair. The matrix ordered according the sign of selection makes unnecessary the comparison of signs of dinucleotides, because this is implicit.

The χ^2^ values of these studies are often enormous and out of tables and programs. I approximated the probability values by the normal or Gaussian equivalent; this is not valid for small degrees of freedom (df), however 9 df may be approximated with a moderate error. The mean and variance of a χ^2^ distribution are the degrees of freedom (df) and twice df, respectively. I made equivalent one point decimal for each two SD from the expected mean (df). With one df the application is no as valid as with nine df, but with enormous figures it yields an approximation to a qualitative intuitive evaluation.

## RESULTS

Table 1 shows the matrix: Selective profiles (columns) vs Separations (Sep) for this strain of the *SARS-CoV-2* virus. Only 4 Seps are in this Table modified from unpublished rusult (C. Y. Valenzuela, submitted for publication). Here, the order of significance include its sign of selection. The most significant pair is CG (-) at Sep 0 with χ^2^_1_ = 377.3, whose probability value is P < 10^-133^.

There are dinucleotides where the Index has the same Par and a-Par dinucleotide: AA, TT, GG and CC whose Par and a-Par are TT, AA, CC and GG, respectively. Other dinucleotides have different Par and equal a-Par as AT, TA, GC and CG whose Par pairs are TA, AT, CG and GC, and whose a-Par pairs are AT, TA, GC, and CG respectively. I analyzed both subsets of pairs only in the comparison of *SARS-CoV-2* with *E. coli* to show a sharp difference in selective profiles with a known big ds DNA genome (see below).

Dinucleotides with different Par and a-Par pairs are AG, AC, TG, TC, GA, GT, CA and CT pairs. I studied this last group of pairs and the first four separations (0, 1, 2 and 3). To explain friendly the method, the result with *SARS-CoV-2* is used. Remember selective profiles include four traits: 1) the order of significance (Sig, 1 to 16) given by 2) the χ^2^_1_ value, 3) the selection coefficient (co-sel), with its 4) sign of selection (+ or -). I compared three of these four traits, but not χ^2^_1_ values between both species due to their difference in genome sizes; the comparison of selective signs is direct and implicit. The difference in selective traits between Par or a-Par and the Index dinucleotides is direct from Table 1. Let us choose TG as the index pair, with 0 Sep, it has Sig = 16° (place), co-sel = +0.377 and Sign = +. Its Par pair is AC with Sig = 14°, co-sel = +0.231 and Sign = +; its a-Par pair is CA with Sig = 15° (place), co-sel = +0.269 and sign = +. The absolute difference in these three traits with the Index is for I-Par pair: 2° Sig, 0.146 co-sel and (+ +) Sign, respectively. The absolute difference in these three traits with the Index is for I-a-Par pair: 1° Sig, 0.108 co-sel and (+ +). Table 2 presents the selective profile comparisons of the Index, Par, and a-Par for the 8 Index dinucleotides having different Par and a-Par pairs, for the four Separations. The Significance distance I-Par was smaller than that of the I-a-Par, but the difference was not significant (minimal significance is P = 0.05). As I mentioned the comparison of signs of selection in the following tables is not included. The differences in the selection coefficient followed the same pattern as Significance, with a higher probability of significance (a less significant result). It is remarkable that this result is different from a previous study (C.Y. Valenzuela, submitted for publication) with distances of the chi-squared value without specifying the sign of selection (OSA).

**Table 2.**
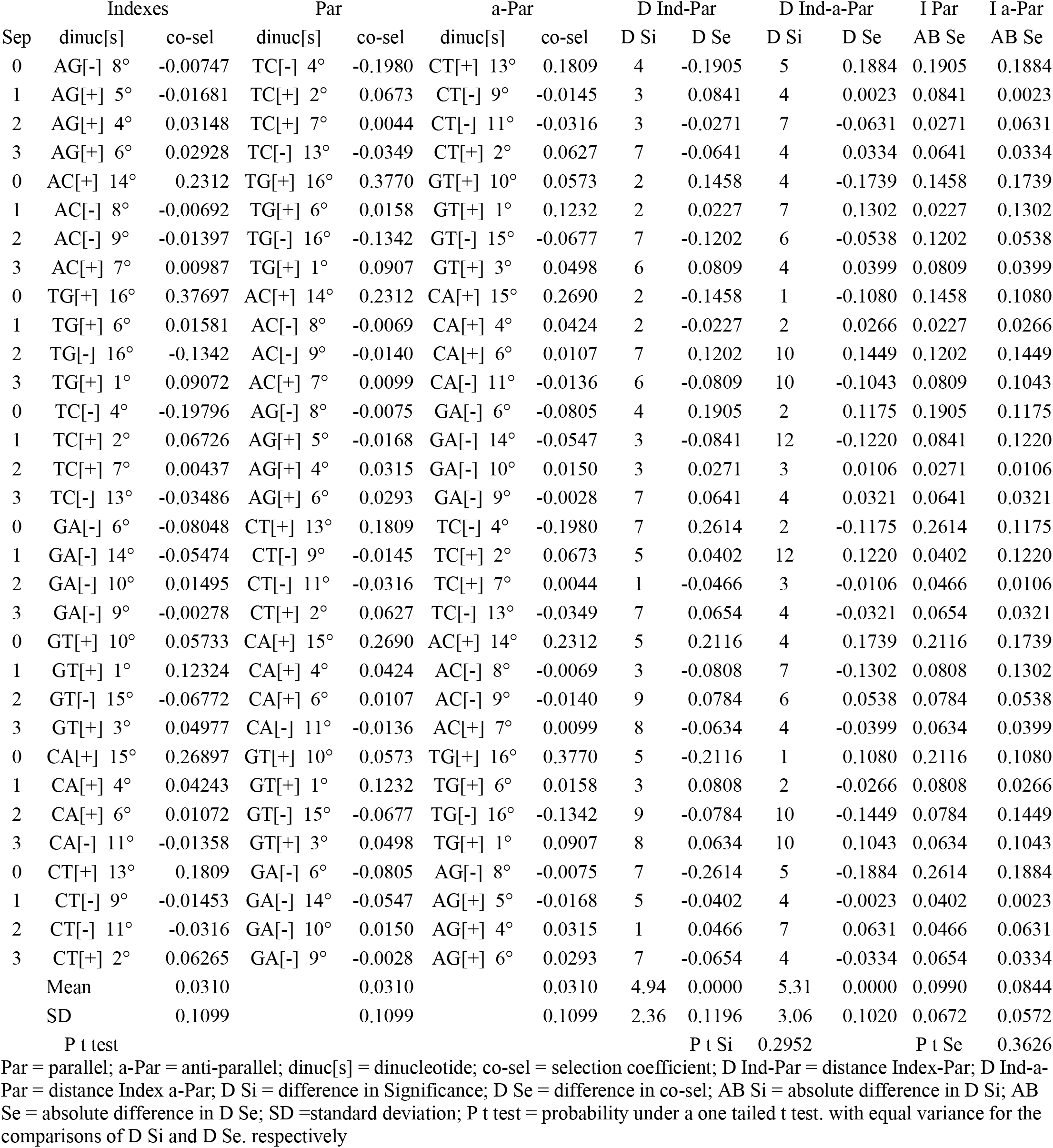
*SARS-CoV-2*. Index dinucleotides with different Par and a-Par pairs. Four Separations.

Before presenting the study of twelve viruses, I present the dinucleotide analysis of the *E. coli* genome to establish clear differences with a rather big (5 million pb) ds DNA. This is necessary because there is not another researcher working in this field and the reader must know what happens with big ds genomes. Table 3 presents the dinucleotide spectrum of the genome analysis of *E coli* with 4 Seps.

**Table 3.** Significance vs Separations of dinucleotides from *E. coli*. Four Separations.

| Sep | dinuc(s) | $\chi^2_1$ | co-sel | dinuc(s) | $\chi^2_1$ | co-sel | dinuc(s) | $\chi^2_1$ | co-sel | dinuc(s) | $\chi^2_1$ | co-sel |
| --- | --- | --- | --- | --- | --- | --- | --- | --- | --- | --- | --- | --- |
|  | 1° Significance |  |  | 2° Significance |  |  | 3° Significance |  |  | 4° Significance |  |  |
| 0 | GC[+] | 24073.2 | 0.2715 | TT[+] | 12324.3 | 0.1998 | AA[+] | 11833.5 | 0.1946 | CG[+] | 6535.8 | 0.1415 |
| 1 | CG[+] | 27665.5 | 0.2911 | AC[+] | 4121.0 | 0.1136 | GT[+] | 3967.0 | 0.1117 | TA[+] | 2204.0 | 0.0842 |
| 2 | GC[-] | 8072.1 | -0.1572 | TG[-] | 6440.1 | -0.1424 | CA[-] | 6434.2 | -0.1420 | GT[-] | 1547.7 | -0.0698 |
| 3 | TA[-] | 7439.1 | -0.1547 | GC[-] | 2552.0 | -0.0884 | AG[-] | 526.3 | -0.0406 | CT[-] | 462.2 | -0.0382 |
|  | 5° Significance |  |  | 6° Significance |  |  | 7° Significance |  |  | 8° Significance |  |  |
| 0 | CA[+] | 4858.4 | 0.1234 | TG[+] | 4746.1 | 0.1222 | AT[+] | 3154.5 | 0.1008 | GA[-] | 1757.2 | -0.0742 |
| 1 | TT[+] | 292.3 | 0.0308 | AA[+] | 279.4 | 0.0299 | GA[+] | 0.4 | 0.0012 | TC[-] | 4.0 | -0.0034 |
| 2 | AC[-] | 1418.3 | -0.0666 | AT[-] | 1245.2 | -0.0633 | TA[-] | 876.4 | -0.0531 | CG[-] | 167.9 | -0.0227 |
| 3 | CG[-] | 195.2 | -0.0245 | CC[-] | 53.4 | -0.0128 | GG[-] | 38.2 | -0.0108 | AT[-] | 5.7 | -0.0043 |
|  | 9° Significance |  |  | 10° Significance |  |  | 11° Significance |  |  | 12° Significance |  |  |
| 0 | TC[-] | 1818.6 | -0.0757 | GG[-] | 2513.4 | -0.0877 | CC[-] | 2557.2 | -0.0885 | AC[-] | 4003.1 | -0.1120 |
| 1 | GC[-] | 83.6 | -0.0160 | AT[-] | 805.5 | -0.0509 | CT[-] | 2682.8 | -0.0919 | CC[-] | 2756.3 | -0.0919 |
| 2 | AG[+] | 245.4 | 0.0277 | CT[+] | 252.7 | 0.0282 | TC[+] | 2621.8 | 0.0909 | GA[+] | 2766.3 | 0.0930 |
| 3 | AA[+] | 13.4 | 0.0066 | TT[+] | 25.4 | 0.0091 | GT[+] | 356.2 | 0.0335 | AC[+] | 470.0 | 0.0384 |
|  | 13° Significance |  |  | 14° Significance |  |  | 15° Significance |  |  | 16° Significance |  |  |
| 0 | GT[-] | 4147.5 | -0.1143 | AG[-] | 9941.2 | -0.1764 | CT[-] | 10128.4 | -0.1786 | TA[-] | 18837.6 | -0.2462 |
| 1 | AG[-] | 2778.5 | -0.0932 | GG[-] | 2872.4 | -0.0937 | TG[-] | 3769.3 | -0.1089 | CA[-] | 4032.0 | -0.1124 |
| 2 | AA[+] | 3295.6 | 0.1027 | TT[+] | 3501.9 | 0.1065 | GG[+] | 5864.6 | 0.1339 | CC[+] | 5873.5 | 0.1342 |
| 3 | TC[+] | 1361.2 | 0.0655 | GA[+] | 1481.5 | 0.0681 | CA[+] | 1844.5 | 0.0760 | TG[+] | 1887.2 | 0.0771 |

Nomenclature as in Table 1

The difference between *SARS-CoV-2* and *E. coli* is evident at the first inspection and comparison between Table 1 and Table 3. Significances in *E. coli* are 50 or more times those of *SARS-CoV-2*. The χ^2^_1_ value of the first dinucleotide GC+ is 24,073.2, this implies a probability (Gaussian approximation; no as valid as in χ^2^_9_ test) less than 10^-4,250^. Dinucleotides with equal Par and a-Par complementary dinucleotides shows sharp differences. In *SARS-CoV-2* AA and TT, and GG and CC separate at the places of significances (Sigs). For example at Sep 0, AA is at 12° Sig while TT is at 9°, the difference in Sig is 3°, at Sep 1 the difference is 7° (3° and 10°, respectively), at Sep 2 is 3° (5° and 2°, respectively), and at Sep 3 it is 3° (12° and 15°, respectively). In *E. coli* AA and TT are contiguous (distance 1°) in the four Seps. GG and CC behave equally, while in *SARS-CoV-2* the distances between both are 2°, 4°, 3° and 4° for Seps 0, 1, 2 and 3, respectively, in *E. coli* they are 1°, 2°, 1° and 1°, respectively. It is evident that in a ds genome *(E. coli*) complementary dinucleotides behave different as in *SARS-CoV-2*, a ss genome, where complementary nucleic acid do not exist. (However, it is remarkable that the difference is not as big as expected; I shall discuss that). This occurs equally with the human genomes (C. Y. Valenzuela, submitted for publication). I remember that the expected mean difference in Sig profile is 5.667 and the standard deviation is 3.636. The following analysis in *E. coli* is similar to that presented in Table 2 for *SARS-CoV-2* on the distance of selective profiles from Par and a-Par dinucleotides to their Index; these Index pairs have different Par and a-Par dinucleotides. Table 4 presents it. In this Table the mean Sig distance I-a-Par is 1.06 (contiguous dinucleotides) while in I-Par is 6.06 (t test P < 10^-9^) and in Sel distance the mean is 0.0679 for I-Par and for I-a-Par is 0.0013 (P < 10^-13^).

**Table 4.** *E. coli*. Index dinucleotides with different Par and a-Par pairs, four Separations.

| Sep | Indexes |  |  | Par |  |  | a-Par |  |  | D Ind - Par |  | D Ind - a-Par |  |
| --- | --- | --- | --- | --- | --- | --- | --- | --- | --- | --- | --- | --- | --- |
|  | dinuc[s] | co-sel | Sig | dinuc[s] | co-sel | Sig | dinuc[s] | co-sel | Sig | D Si | D Se | D Si | D Se |
| 1 | AG[-] | -0.1764 | 14 | TC[-] | -0.0757 | 9 | CT[-] | -0.1786 | 15 | 5 | 0.1007 | 1 | 0.0023 |
| 2 | AG[-] | -0.0932 | 13 | TC[-] | -0.0034 | 8 | CT[-] | -0.0919 | 11 | 5 | 0.0899 | 2 | 0.0013 |
| 3 | AG[+] | 0.0277 | 9 | TC[+] | 0.0909 | 11 | CT[+] | 0.0282 | 10 | 2 | 0.0632 | 1 | 0.0005 |
| 4 | AG[-] | -0.0406 | 3 | TC[+] | 0.0655 | 13 | CT[-] | -0.0382 | 4 | 10 | 0.1061 | 1 | 0.0024 |
| 1 | AC[-] | -0.1120 | 12 | TG[+] | 0.1222 | 6 | GT[-] | -0.1143 | 13 | 6 | 0.2342 | 1 | 0.0023 |
| 2 | AC[+] | 0.1136 | 2 | TG[-] | -0.1089 | 15 | GT[+] | 0.1117 | 3 | 13 | 0.2225 | 1 | 0.0019 |
| 3 | AC[-] | -0.0666 | 5 | TG[-] | -0.1424 | 2 | GT[-] | -0.0698 | 4 | 3 | 0.0757 | 1 | 0.0032 |
| 4 | AC[+] | 0.0384 | 12 | TG[+] | 0.0771 | 16 | GT[+] | 0.0335 | 11 | 4 | 0.0387 | 1 | 0.0049 |
| 1 | TG[+] | 0.1222 | 6 | AC[-] | -0.1120 | 12 | CA[+] | 0.1234 | 5 | 6 | 0.2342 | 1 | 0.0011 |
| 2 | TG[-] | -0.1089 | 15 | AC[+] | 0.1136 | 2 | CA[-] | -0.1124 | 16 | 13 | 0.2225 | 1 | 0.0034 |
| 3 | TG[-] | -0.1424 | 2 | AC[-] | -0.0666 | 5 | CA[-] | -0.1420 | 3 | 3 | 0.0757 | 1 | 0.0004 |
| 4 | TG[+] | 0.0771 | 16 | AC[+] | 0.0384 | 12 | CA[+] | 0.0760 | 15 | 4 | 0.0387 | 1 | 0.0011 |
| 1 | TC[-] | -0.0757 | 9 | AG[-] | -0.1764 | 14 | GA[-] | -0.0742 | 8 | 5 | 0.1007 | 1 | 0.0015 |
| 2 | TC[-] | -0.0034 | 8 | AG[-] | -0.0932 | 13 | GA[+] | 0.0012 | 7 | 5 | 0.0899 | 1 | 0.0045 |
| 3 | TC[+] | 0.0909 | 11 | AG[+] | 0.0277 | 9 | GA[+] | 0.0930 | 12 | 2 | 0.0632 | 1 | 0.0022 |
| 4 | TC[+] | 0.0655 | 13 | AG[-] | -0.0406 | 3 | GA[+] | 0.0681 | 14 | 10 | 0.1061 | 1 | 0.0026 |
| 1 | GA[-] | -0.0742 | 8 | CT[-] | -0.1786 | 15 | TC[-] | -0.0757 | 9 | 7 | 0.1045 | 1 | 0.0015 |
| 2 | GA[+] | 0.0012 | 7 | CT[-] | -0.0919 | 11 | TC[-] | -0.0034 | 8 | 4 | 0.0931 | 1 | 0.0045 |
| 3 | GA[+] | 0.0930 | 12 | CT[+] | 0.0282 | 10 | TC[+] | 0.0909 | 11 | 2 | 0.0648 | 1 | 0.0022 |
| 4 | GA[+] | 0.0681 | 14 | CT[-] | -0.0382 | 4 | TC[+] | 0.0655 | 13 | 10 | 0.1062 | 1 | 0.0026 |
| 1 | GT[-] | -0.1143 | 13 | CA[+] | 0.1234 | 5 | AC[-] | -0.1120 | 12 | 8 | 0.2376 | 1 | 0.0023 |
| 2 | GT[+] | 0.1117 | 3 | CA[-] | -0.1124 | 16 | AC[+] | 0.1136 | 2 | 13 | 0.2241 | 1 | 0.0019 |
| 3 | GT[-] | -0.0698 | 4 | CA[-] | -0.1420 | 3 | AC[-] | -0.0666 | 5 | 1 | 0.0722 | 1 | 0.0032 |
| 4 | GT[+] | 0.0335 | 11 | CA[+] | 0.0760 | 15 | AC[+] | 0.0384 | 12 | 4 | 0.0425 | 1 | 0.0049 |
| 1 | CA[+] | 0.1234 | 5 | GT[-] | -0.1143 | 13 | TG[+] | 0.1222 | 6 | 8 | 0.2376 | 1 | 0.0011 |
| 2 | CA[-] | -0.1124 | 16 | GT[+] | 0.1117 | 3 | TG[-] | -0.1089 | 15 | 13 | 0.2241 | 1 | 0.0034 |
| 3 | CA[-] | -0.1420 | 3 | GT[-] | -0.0698 | 4 | TG[-] | -0.1424 | 2 | 1 | 0.0722 | 1 | 0.0004 |
| 4 | CA[+] | 0.0760 | 15 | GT[+] | 0.0335 | 11 | TG[+] | 0.0771 | 16 | 4 | 0.0425 | 1 | 0.0011 |
| 1 | CT[-] | -0.1786 | 15 | GA[-] | -0.0742 | 8 | AG[-] | -0.1764 | 14 | 7 | 0.1045 | 1 | 0.0023 |
| 2 | CT[-] | -0.0919 | 11 | GA[+] | 0.0012 | 7 | AG[-] | -0.0932 | 13 | 4 | 0.0931 | 2 | 0.0013 |
| 3 | CT[+] | 0.0282 | 10 | GA[+] | 0.0930 | 12 | AG[+] | 0.0277 | 9 | 2 | 0.0648 | 1 | 0.0005 |
| 4 | CT[-] | -0.0382 | 4 | GA[+] | 0.0681 | 14 | AG[-] | -0.0406 | 3 | 10 | 0.1062 | 1 | 0.0024 |
| M |  | -0.0178 | 9.41 |  | -0.0178 | 4.41 |  | -0.0178 | 9.41 | 6.06 | 0.1172 | 1.06 | 0.0022 |
| SD |  | 0.0938 | 4.48 |  | 0.0938 | 4.48 |  | 0.0938 | 4.48 | 3.67 | 0.0679 | 0.24 | 0.0013 |
| P t |  |  |  |  |  |  |  |  |  |  |  | 1.1E-10 | 7E-14 |

Nomenclature as in Table 2. D Ind-Par = absolute distance Index-Par; D Ind -a-Par = absolute distance Index-a-Par

Selective profile of the 5’-3’ a-Par dinucleotide is practically the same as the selective profile of the Index. The 5’-3’ parallel dinucleotide differs from the Index. Remark that it is not possible to analyze the true 3’-5’ parallel dinucleotide, because we have only one strand from GenBank. The correction of this possible error is, at present OSA. Statistical tests where done as compliance of statistical norms, because values of differences in Sigs and co-sel are significant by the simple inspection. Differences in co-sel in the I-Par comparisons are at the order of tenths or hundredths, and the differences in I-a-Par are at the order of thousandths or ten thousandths. In Table 4 only absolute distances I-Par and I-a-Par are considered.

Table 5 for DNA and Table 6 for RNA viruses present the comparisons for 12 viruses with the same format of *SARS-CoV-2* and *E. coli*. Only mean, SD and t test probability appear. The single or double stranded DNA or RNA condition is in the first row of each set of viruses. Names are those used in current language and in GenBank and not necessarily scientific species nomenclature (excepting *SARS-CoV-2* and *E.coli*, see accession numbers of genomes).

**Table 5.** Means, standard deviations and t test probability for distances Index-Par and Index-a-Par. DNA viruses.

|  | Indexes<br>co-sel | Par<br>co-sel | a-Par<br>co-sel | D Ind - Par |  | D Ind - a-Par |  | Ind Par<br>AB Se | Ind a-Par<br>AB Se |
| --- | --- | --- | --- | --- | --- | --- | --- | --- | --- |
|  |  |  |  | D Si | D Se | D Si | D Se |  |  |
| Double stranded DNA viruses |  |  |  |  |  |  |  |  |  |
| Human adenovirus C strain KF268127.1; 35,931 bp |  |  |  |  |  |  |  |  |  |
| Mean | -0.0414 | -0.0414 | -0.0414 | 4.7188 | 0.0000 | 3.5000 | 0.0000 | 0.0752 | 0.0417 |
| SD | 0.0649 | 0.0649 | 0.0649 | 2.2533 | 0.1002 | 1.7854 | 0.0456 | 0.0662 | 0.0184 |
| Prob t |  |  |  |  | P t Si = | 0.0107 |  | P t Se = | 0.0043 |
| Human cytomegalovirus; 229,354 bp |  |  |  |  |  |  |  |  |  |
| Mean | -0.0304 | -0.0304 | -0.0304 | 4.1875 | 0.0000 | 1.1875 | 0.0000 | 0.0491 | 0.0088 |
| SD | 0.0580 | 0.0580 | 0.0580 | 2.7207 | 0.0582 | 0.3903 | 0.0127 | 0.0312 | 0.0091 |
| Prob t |  |  |  |  | P t Si = | 4.1E-08 |  | P t Se = | 1.7E-09 |
| Escherichia phage 2; 167,318 bp |  |  |  |  |  |  |  |  |  |
| Mean | 0.0089 | 0.0089 | 0.0089 | 4.75 | 0.0000 | 2.625 | 0.0000 | 0.0555 | 0.0346 |
| SD | 0.0554 | 0.0554 | 0.0554 | 3.0923 | 0.0745 | 1.9325 | 0.0512 | 0.0498 | 0.0378 |
| Prob t |  |  |  |  | P t Si = | 0.0009 |  | P t Se = | 0.0339 |
| His2 virus; 16,067 bp |  |  |  |  |  |  |  |  |  |
| Mean | 0.0500 | 0.0500 | 0.0500 | 3.5313 | 0.0000 | 1.5313 | 0.0000 | 0.07576 | 0.0368 |
| SD | 0.1099 | 0.1099 | 0.1099 | 2.1935 | 0.1026 | 0.8286 | 0.0441 | 0.0692 | 0.0244 |
| Prob t |  |  |  |  | P t Si = | 6.3E-6 |  | P t Se = | 0.0022 |
| Single stranded DNA Viruses |  |  |  |  |  |  |  |  |  |
| Penaeus monodon hepadensovirus 4; 6,310 b |  |  |  |  |  |  |  |  |  |
| Mean | 0.0186 | 0.0186 | 0.0186 | 5.2500 | 0.0000 | 4.8750 | 0.0000 | 0.1034 | 0.1016 |
| SD | 0.1034 | 0.1034 | 0.1034 | 3.4369 | 0.1507 | 3.2763 | 0.1341 | 0.1096 | 0.0875 |
| Prob t |  |  |  |  | P t Si = | 0.3308 |  | P t Se = | 0.4731 |
| Escherichia phage phi-X174; 5,386 b |  |  |  |  |  |  |  |  |  |
| Mean | 0.0038 | 0.0038 | 0.0038 | 3.4375 | 0.0000 | 7.125 | 0.0000 | 0.1038 | 0.1721 |
| SD | 0.1174 | 0.1174 | 0.1174 | 2.0606 | 0.1343 | 2.966 | 0.2016 | 0.0852 | 0.1051 |
| Prob t |  |  |  |  | P t Si = | 1.9E-07 |  | P t Se = | 0.0033 |
| Phage M13 Enterobacteria; 6,407 b |  |  |  |  |  |  |  |  |  |
| Mean | -0.0271 | -0.0271 | -0.0271 | 4.3125 | 0.0000 | 5.9375 | 0.0000 | 0.0952 | 0.1062 |
| SD | 0.0939 | 0.0939 | 0.0939 | 3.0765 | 0.1309 | 3.7992 | 0.1442 | 0.0899 | 0.0975 |
| Prob t |  |  |  |  | P t Si = | 0.0345 |  | P t Se = | 0.3234 |

Nomenclature as in Table 2

**Table 6.**
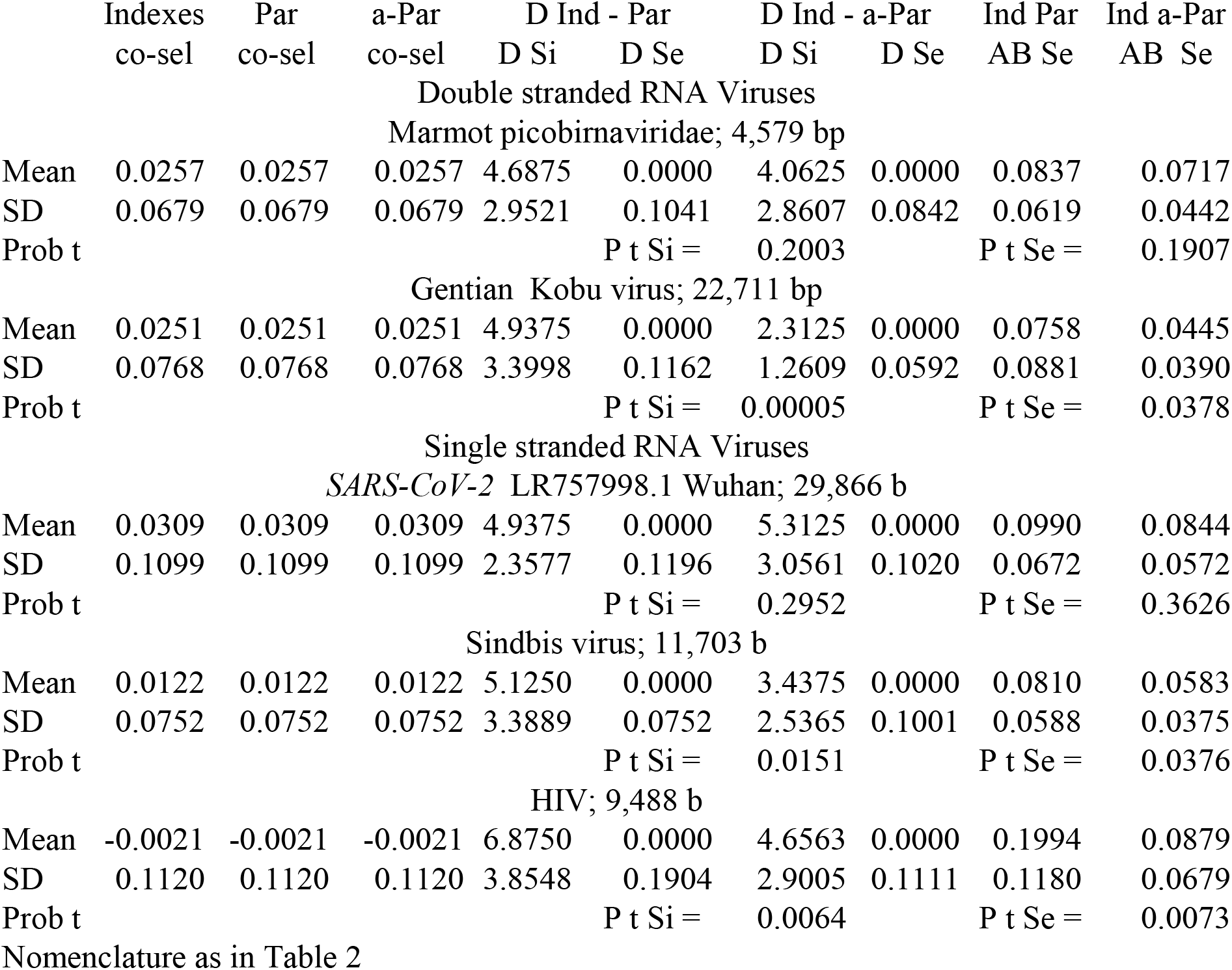
Means. standard deviations and t test probability for distances Index-Par and Index-a-Par. RNA viruses.

|  | Indexes | Par | a-Par | D Ind - Par |  | D Ind - a-Par |  | Ind Par | Ind a-Par |
| --- | --- | --- | --- | --- | --- | --- | --- | --- | --- |
|  | co-sel | co-sel | co-sel | D Si | D Se | D Si | D Se | AB Se | AB Se |
| Double stranded RNA Viruses |  |  |  |  |  |  |  |  |  |
| Marmot picobirnaviridae; 4,579 bp |  |  |  |  |  |  |  |  |  |
| Mean | 0.0257 | 0.0257 | 0.0257 | 4.6875 | 0.0000 | 4.0625 | 0.0000 | 0.0837 | 0.0717 |
| SD | 0.0679 | 0.0679 | 0.0679 | 2.9521 | 0.1041 | 2.8607 | 0.0842 | 0.0619 | 0.0442 |
| Prob t |  |  |  | P t Si = |  | 0.2003 |  | P t Se = | 0.1907 |
| Gentian Kobu virus; 22,711 bp |  |  |  |  |  |  |  |  |  |
| Mean | 0.0251 | 0.0251 | 0.0251 | 4.9375 | 0.0000 | 2.3125 | 0.0000 | 0.0758 | 0.0445 |
| SD | 0.0768 | 0.0768 | 0.0768 | 3.3998 | 0.1162 | 1.2609 | 0.0592 | 0.0881 | 0.0390 |
| Prob t |  |  |  | P t Si = |  | 0.00005 |  | P t Se = | 0.0378 |
| Single stranded RNA Viruses |  |  |  |  |  |  |  |  |  |
| SARS-CoV-2 LR757998.1 Wuhan; 29,866 b |  |  |  |  |  |  |  |  |  |
| Mean | 0.0309 | 0.0309 | 0.0309 | 4.9375 | 0.0000 | 5.3125 | 0.0000 | 0.0990 | 0.0844 |
| SD | 0.1099 | 0.1099 | 0.1099 | 2.3577 | 0.1196 | 3.0561 | 0.1020 | 0.0672 | 0.0572 |
| Prob t |  |  |  | P t Si = |  | 0.2952 |  | P t Se = | 0.3626 |
| Sindbis virus; 11,703 b |  |  |  |  |  |  |  |  |  |
| Mean | 0.0122 | 0.0122 | 0.0122 | 5.1250 | 0.0000 | 3.4375 | 0.0000 | 0.0810 | 0.0583 |
| SD | 0.0752 | 0.0752 | 0.0752 | 3.3889 | 0.0752 | 2.5365 | 0.1001 | 0.0588 | 0.0375 |
| Prob t |  |  |  | P t Si = |  | 0.0151 |  | P t Se = | 0.0376 |
| HIV; 9,488 b |  |  |  |  |  |  |  |  |  |
| Mean | -0.0021 | -0.0021 | -0.0021 | 6.8750 | 0.0000 | 4.6563 | 0.0000 | 0.1994 | 0.0879 |
| SD | 0.1120 | 0.1120 | 0.1120 | 3.8548 | 0.1904 | 2.9005 | 0.1111 | 0.1180 | 0.0679 |
| Prob t |  |  |  | P t Si = |  | 0.0064 |  | P t Se = | 0.0073 |
| Nomenclature as in Table 2 |  |  |  |  |  |  |  |  |  |

Nomenclature as in Table 2

These results were expected; ds DNA viruses showed differences between I-Par larger than differences between I-a-Par. Cytomegalovirus had the largest difference very probably because it is the largest virus. On the contrary, ss DNA viruses disagreed with this tendency. Penaeus monodon did not present significant difference between I-Par and I-a-Par differences. However and unexpectedly Coliphage phi-X174 presented an inverse difference of the differences: I-Par difference is significant smaller than the I-a-Par difference.

Table 6 shows RNA viruses. In ds RNA viruses Marmot picobirnaviridae virus presented the expected larger difference in I-Par pair, but no significantly, while Gentian Kobu virus did. In this case, the small size of the viruses may explain the non-significant result. As expected the ss RNA virus *SARS-CoV-2* did not presented difference between I-Par and I-a-Par, while Sindbis virus presented a small but significant difference and HIV virus a large and significant difference.

I mentioned these distances to neutral evolution were periodical. It seems important to examine whether these viromes present periodicity because this is the first time the method applies to a set of viruses*. SARS-CoV-2* is periodical (C. Y. Valenzuela. submitted for publication) the 12 viromes and *E coli* are in Table 7. The pivot distance (the largest) of the period is in bold character it follows and precedes smaller values. Underlined values of the series do not accomplish partially or completely this criterion. Those values less than 17 are not significant and do not alter necessarily the periodicity in this set of values. This is because the variance of the test is 2xdf = 18.0 and the SD is 4.243; random fluctuation near those values blurs the periodicity. Besides the χ^2^_9_ value is less significant in ss DNA or RNA and smaller viruses. Regardless these restrictions the periodicity is impressive.

**Table 7.** Periodicities of distances to evolutionary neutrality in viromes and *E. coli* genome.

| Sep | Adenov | Cytome | Ph 2 | His2 | phiX | hepand | ph M13 | marmot | Gentian | SARS | Sindbis | HIV | *E. coli* |
| --- | --- | --- | --- | --- | --- | --- | --- | --- | --- | --- | --- | --- | --- |
| 0 | 509.1 | 2981.2 | 1237.4 | 306.6 | 132.0 | 196.2 | 144.9 | 41.5 | 684.3 | 1236.9 | 107.4 | 464.5 | 123230.0 |
| 1 | 140.3 | 1581.4 | 888.9 | 318.1 | 92.8 | 46.9 | 61.9 | 32.3 | 83.3 | 75.7 | 95.8 | 83.7 | 58312.6 |
| 2 | **424.2** | **3808.1** | **488.1** | **150.6** | **61.0** | **33.6** | **33.2** | **26.7** | **76.1** | **144.0** | **55.2** | **61.3** | **50623.7** |
| 3 | 44.3 | 379.9 | 432.3 | 73.5 | 53.6 | 10.9 | 24.0 | 10.8 | 38.7 | 69.5 | 24.9 | 22.4 | 18711.1 |
| 4 | 13.4 | 308.8 | 254.3 | 66.6 | 25.8 | 18.7 | 20.9 | 11.9 | 14.4 | 88.8 | 5.8 | 34.7 | 5443.7 |
| 5 | **413.2** | **2660.8** | **392.0** | <u>**59.3**</u> | **47.6** | <u>**18.4**</u> | **32.3** | **22.2** | **24.7** | <u>**72.1**</u> | <u>**15.7**</u> | **23.0** | **8927.9** |
| 6 | 22.0 | 112.8 | 292.8 | 46.0 | 18.7 | 8.4 | 20.4 | 5.4 | 18.7 | 72.7 | 16.2 | 31.2 | 1363.4 |
| 7 | 23.6 | 61.6 | 247.3 | 78.9 | 29.9 | 17.7 | 11.4 | 7.8 | 8.2 | 44.9 | 16.3 | 11.4 | 2052.5 |
| 8 | **356.3** | **2419.4** | **523.6** | **120.0** | **87.2** | **30.8** | **50.8** | **8.5** | **42.9** | **127.6** | <u>**12.0**</u> | **45.9** | **10597.0** |
| 9 | 13.4 | 47.1 | 320.6 | 33.4 | 28.3 | 12.6 | 20.9 | 12.1 | 15.6 | 56.0 | 3.3 | 10.0 | 2138.6 |
| 10 | 9.3 | 112.0 | 309.2 | 17.3 | 40.0 | 13.7 | 26.5 | 15.5 | 29.4 | 40.4 | 14.7 | 18.9 | 2346.0 |
| 11 | **311.0** | **2530.4** | **583.1** | **95.1** | **68.4** | **29.7** | **60.6** | **20.6** | **53.2** | **131.6** | **24.3** | **47.2** | **16223.7** |
| 12 | 25.8 | 107.7 | 243.6 | 44.5 | 44.8 | 27.0 | 32.4 | 18.5 | 40.8 | 85.8 | 18.1 | 35.2 | 1986.7 |
| 13 | 13.5 | 51.7 | 213.1 | 36.4 | 28.1 | 6.1 | 26.1 | 13.1 | 12.4 | 70.3 | 19.8 | 15.1 | 1676.3 |
| 14 | **278.7** | **1978.6** | **468.1** | **64.6** | **47.9** | **32.8** | **72.0** | **18.0** | <u>**9.4**</u> | **101.1** | <u>**13.4**</u> | **23.9** | **9563.7** |
| 15 | 11.8 | 69.4 | 157.9 | 15.4 | 38.4 | 7.7 | 28.5 | 14.9 | 29.9 | 57.1 | 7.2 | 14.8 | 1647.4 |
| 16 | 18.6 | 88.9 | 200.7 | 25.5 | 46.9 | 15.5 | 35.2 | 9.0 | 25.9 | 61.4 | 31.7 | 17.2 | 2235.3 |
| 17 | **287.6** | **1989.8** | **423.0** | **54.0** | **49.6** | **22.6** | **52.4** | **12.4** | **42.7** | **86.4** | <u>**18.1**</u> | **36.8** | **9725.6** |
| 18 | 3.4 | 38.2 | 155.4 | 16.8 | 25.7 | 9.2 | 39.9 | 7.1 | 15.5 | 46.5 | 9.3 | 10.0 | 1566.9 |
| 19 | 18.3 | 74.4 | 176.4 | 25.3 | 26.8 | 11.8 | 44.1 | 9.7 | 13.4 | 57.3 | 12.5 | 28.0 | 1940.5 |
| 20 | **254.3** | **1935.0** | **534.5** | **70.3** | **69.6** | **34.8** | **54.7** | **20.8** | **30.0** | **93.9** | **21.8** | **34.8** | **13582.1** |
| 21 | 24.4 | 82.5 | 183.1 | 28.9 | 20.7 | 17.4 | 20.8 | 18.2 | 27.5 | 87.4 | 11.2 | 7.4 | 2220.1 |
| 22 | 12.0 | 61.3 | 235.3 | 23.0 | 29.2 | 14.4 | 31.8 | 30.7 | 26.4 | 63.3 | 20.8 | 18.0 | 1661.0 |
| 23 | **225.4** | **1821.7** | **500.8** | **78.9** | **42.6** | **39.3** | **61.9** | <u>**14.8**</u> | <u>**25.6**</u> | **124.8** | **31.3** | <u>**15.6**</u> | **12420.6** |
| 24 | 22.3 | 79.9 | 153.9 | 9.4 | 27.0 | 16.6 | 37.1 | 17.0 | 16.4 | 67.8 | 15.9 | 17.7 | 2320.9 |
| 25 | 15.1 | 57.1 | 190.1 | 12.5 | 31.3 | 19.6 | 20.4 | 7.3 | 17.1 | 54.7 | 20.9 | 8.9 | 1684.5 |
| 26 | **234.6** | **1651.8** | **482.0** | **68.0** | **42.6** | **37.9** | **58.6** | **25.0** | <u>**20.2**</u> | **74.1** | <u>**18.7**</u> | **18.4** | **10400.1** |
| 27 | 24.2 | 79.8 | 209.0 | 10.7 | 20.9 | 11.8 | 28.7 | 17.1 | 27.5 | 59.1 | 6.7 | 13.6 | 2078.5 |
| 28 | 22.4 | 51.8 | 240.3 | 14.6 | 27.0 | 11.3 | 20.3 | 18.1 | 5.0 | 45.0 | 15.5 | 9.1 | 1926.3 |
| 29 | **189.4** | **1805.1** | **456.9** | **72.5** | **59.4** | <u>**10.6**</u> | **55.1** | <u>**12.5**</u> | **34.2** | **64.2** | **18.5** | **22.4** | **10553.1** |
| 30 | 23.8 | 45.4 | 206.0 | 25.6 | 28.1 | 13.4 | 17.1 | 13.6 | 4.7 | 42.5 | 9.7 | 11.2 | 2215.2 |
| 31 | 15.7 | 44.4 | 180.8 | 19.9 | 27.6 | 17.4 | 14.4 | 15.9 | 14.1 | 40.2 | 6.9 | 7.0 | 1817.7 |
| 32 | **172.3** | **1593.0** | **534.8** | **62.2** | **65.0** | <u>**15.8**</u> | **32.3** | **42.6** | **35.4** | **124.9** | **11.3** | **44.3** | **12264.9** |

In bold type the largest distance among the triplet of the period measured by the total $\chi^2_9$ . For names of viruses see Genomes.

Moreover, we see that single or double stranded viruses that may incorporate to the host dsDNA present a clear periodicity similar as the host one without exceptions (HIV presents only one exception at Sep 23).

## DISCUSSION

This is the first time this statistical tool is used to search for the single or double stranded nature of RNA and DNA viruses and their implications on evolution. Results are according to the theoretical expected behavior of ds or ss viruses with some important exceptions. Escherichia phage phi X174, an ss DNA virus presented a more similar selective profile between Indices and Par pairs than between Indices and a-Par pairs. A plausible explanation is the fact that this virus replicates by using a negative DNA strand after constructing a dsDNA (6, 7). The Marmot Picobirnaviridae a dsRNA virus (8) shows a non-significant larger selective profile difference between I-Par than I-a-Par, this is perhaps it is a very small virus (4.579 bp). However, Gentian Kobu sho-associated virus also a dsRNA virus (9) according to the Oxford database shows a big significant difference as expected; however, other authors present it as an ssRNA virus (10). According to my analysis, this virus behaves as an evolutionary or replicative dsRNA virus. Arguments in favor or against are present in these analyses and Tables but their study is OSA. In the group of ssRNA, viruses *SARS-CoV-2* did not presented differences between I-Par and I-a-Par pairs selective profiles, while Sindbis virus and HIV showed a significant difference where I-Par differences were larger than I-a-Par differences. This was more significant in HIV than in Sindbis virus perhaps for HIV is a retrovirus that exists as a double stranded DNA segment incorporated in the host DNA; however, Sindbis virus needs ds RNA phase in its replication cycle (11). These data, methods and analyses were presented in international and national congresses (12, 13).

The present analyses indicate that the classification in single or double stranded DNA or RNA virus based on the virion physical constitution is useful but it does not include evolutionarily functional double or single stranded constitution mostly produced at the replicative or transcription stage of the viral vital cycle. This new conceptualization seems useful and offers a different understanding of the viral behavior and evolution, which is also useful in epidemiological studies. However, to analyze more precisely these hypotheses need a deep research (OSA).

As for the impressive periodicities, two hypotheses may be advanced. The structural hypothesis proposes these periodicities of the distance to neutrality originated in a basic property of nucleic acids that has different structural stability in triplets of nucleotides; triplets for the genetic code are a result of adaptation to this primary periodicity. The functional hypothesis proposes the periodicity has been acquired as the genetic code installed. During and after this installation the genome segments (coding or non-coding for proteins) optimize an organization that dances at the rhythm of trinucleotides. Arguments in favor or against of these hypotheses are OSA. I would like to remark that the comparisons of differences in selective profiles between an index (I) dinucleotide with its complementary parallel (Par) and anti-parallel (a-Par) dinucleotide informs on the double or single stranded (ds or ss) nature of the nucleic acid of any organism. The foundation of this discrimination happens because ds DNA or ds RNA have parallel and antiparallel strands that evolve together, but ss DNA or ss RNA do not. The single and double stranded evolutionary functional constitution of viruses appears as a useful tool to understand the life cycle of viruses and their epidemiological properties.

## ACKNOWLEDGEMENTS

To Francesco M Scudo and Ching Chung Li my first teachers of population genetics. My students Franccesca Soriano, Francisco Rosas and Matías Rozas helped me in processing data.

## AUTHOR AFFILIATION

Programa de Genética Humana. Instituto de Ciencias Biomédicas (ICBM). Facultad de Medicina. Universidad de Chile. Santiago. Chile

**AUTHOR ORCID** ID 0000-0002-2235-1050

## AUTHOR’S CONTRIBUTION

CYV carried out all the work described in this article

## AUTHOR’S INFORMATION

**Full address and affiliation**: Carlos Y Valenzuela, Programa de Genética Humana. Instituto de Ciencias Biomédicas (ICBM), Facultad de Medicina, Universidad de Chile. Independencia 1027, Código Postal 8380453, Independencia, CHILE. FAX (56–2) 27373158; Phone (56–2) 29786302. E.

## COMPETING INTERESTS

I do not have any competing interest.

## AVAILABILITY OF DATA AND MATERIAL

The genomes’ information is available in GenBank as cited. Any geneticist, student or statistician who knows the current informatics languages can construct these computer programs and perform these analyses.

## CONSENT FOR PUBLICATION

Not applicable.

## ETHICS APPROVAL AND CONSENT TO PARTICIPATE

Not applicable.

## FUNDING

This work and the article did not have extraordinary funds.

